# Integrative Transcriptomic Meta-Analysis Reveals SIRT1-SNCA Correlation in Parkinson’s Disease

**DOI:** 10.64898/2026.11.03.763624

**Authors:** Fatma Hotaslier, Ekin Sonmez, Huseyin Cimen

**Affiliations:** Department of Molecular Biology and Genetics, Gebze Technical University, Kocaeli, Turkiye; The Institute of Biotechnology, Gebze Technical University, Kocaeli, Turkiye; Central Research Laboratory Application and Research Center (GTUMAR), Gebze Technical University, Kocaeli, Turkiye

**Keywords:** Parkinson’s disease, resveratrol, SIRT1, α-synuclein, drug repurposing, transcriptomic meta-analysis

## Abstract

Parkinson’s disease (PD) is a progressive neurodegenerative disorder that affects midbrain dopaminergic neurons without any approved therapy. Candidate molecular targets identified in one model are not always traceable across the postmortem, induced pluripotent stem cell (iPSC), and cell-based systems used to study PD, which complicates clinical translation. To address this gap, we performed an integrative transcriptomic meta-analysis of 33 publicly available datasets across postmortem human brain, iPSC-derived models, and dopaminergic cell lines of PD. Differential expression and functional enrichment analyses were combined with STRING protein-protein interaction networks, followed by drug-gene interaction screening against a polyphenol panel. Overall, resveratrol was recovered in all 33 comparisons, independent of model system, tissue, and genetic background. We associated resveratrol with 30 distinct target genes, and SIRT1 was itself a recovered target in 6 comparisons spanning all three model systems. Finally, we resolved the SIRT1-SNCA relationship into four directional patterns that differed by model, inverse in postmortem brain and positive in acutely toxin-exposed cells. These findings support a candidate, context-dependent resveratrol-SIRT1 axis in PD.

## 1 Introduction

Parkinson’s disease (PD) is the second most common neurodegenerative disorder after Alzheimer’s disease, affecting 2–3% of the population aged 65 years and older (1). Pathologically, it is defined by the progressive degeneration of dopaminergic neurons in the substantia nigra and by Lewy bodies formed through abnormal α-synuclein aggregation (2,3). Dopaminergic cell death has been linked to mitochondrial dysfunction, disrupted dopamine metabolism, oxidative stress, neuroinflammation, and ubiquitin-proteasome system impairment. This molecular complexity parallels the clinical heterogeneity of PD, which spans motor features such as bradykinesia, tremor, and rigidity as well as cognitive, autonomic, and affective symptoms, and indicates that PD is a syndrome comprising distinct phenotypes rather than a uniform disorder without clinical therapy (4, 5).

The etiology remains unknown in most patients. Genetic variants (5–15% of cases), environmental exposures, and their interaction are all implicated (6, 7). Mutations in *SNCA*, *LRRK2*, *PRKN,* and *PINK1*, and *GBA1* risk variants have shaped current models of pathogenesis. *SNCA*, the first gene linked to hereditary PD, encodes α-synuclein and is central to both familial and sporadic disease. It is the main component of Lewy bodies, and its mutations are associated with earlier onset, faster motor progression, and accelerated cognitive decline (3, 5, 8). Environmental factors, including pesticides (rotenone and paraquat), heavy metals, and neurotoxins (MPTP/MPP⁺ and 6-OHDA), increase the risk of PD through mitochondrial dysfunction, oxidative stress, and neuroinflammation (9).

Given the multifactorial and region-specific nature of PD pathology, its molecular mechanisms have been investigated using complementary platforms, including postmortem human brain tissue, induced pluripotent stem cell (iPSC)-derived models, and cell-based systems (10, 14, 16). Postmortem tissue provides the only direct access to the diseased human brain and to the regional progression of Lewy pathology across Braak stages (10, 11). However, transcriptomic analyses of the *substantia nigra* are complicated by the 60–80% dopaminergic neuron loss already present at diagnosis, limited tissue availability, postmortem delay, and inter-individual heterogeneity (10–12), which can be partly addressed by examining regions affected at different disease stages, such as the frontal cortex, ventral tegmental area, and putamen (13). The technology of iPSC enables the generation of patient-specific dopaminergic neurons from individuals with idiopathic PD or corresponding mutations, providing a platform for mechanistic studies and drug screening. However, donor-to-donor variability and the absence of native tissue architecture remain limitations (14, 15). Cell-based models, including SH-SY5Y and SK-N-MC neuroblastoma cells and mesencephalic LUHMES cells, offer scalable systems in which these neurotoxins reproduce key features of dopaminergic neurodegeneration (16); among them, LUHMES cells show the most robust dopaminergic phenotype and the greatest toxin sensitivity (17). Each system therefore captures a different facet of PD as the end-stage pathology in the human brain, patient genetic background in iPSC-derived neurons, and acute, controlled toxic insult in cell lines. Because the systems also differ in platform, cell composition, and disease stage, pooling them would confound biological signal with model-specific variation. Treating them as three parallel groups analyzed with identical thresholds means that a signal recovered independently in all three is unlikely to reflect the idiosyncrasies of any single model (18).

Despite these differences, mitochondrial dysfunction and oxidative stress are reproduced in all three systems, from complex I deficiency in postmortem brain to mitochondrial defects in genetic models and toxin-induced respiratory failure in cell lines, which makes them core and therapeutically tractable processes (9, 16, 19). Within this framework, the NAD⁺-dependent deacetylase SIRT1 acts as a regulatory hub, and it supports mitochondrial biogenesis and antioxidant defense through the FOXO and PGC-1α pathways, limits cellular damage by modulating p53 activity and suppressing NF-κB and PARP-1 signaling, and promotes autophagy and mitophagy (20). Autophagic activation in turn facilitates the clearance of misfolded proteins, including α-synuclein, directly linking SIRT1 activity to *SNCA* accumulation and positioning SIRT1 at the intersection of the two principal axes of PD pathology (21).

Resveratrol, a natural stilbene found in grapes, berries, and peanuts, is the most widely studied SIRT1-activating compound, and it reduces mitochondrial dysfunction, oxidative stress, and α-synuclein accumulation in PD models (22, 23). Clinical evidence comes mainly from Alzheimer’s disease, where a phase 2 trial found resveratrol safe and well tolerated and detected it in cerebrospinal fluid, confirming that it reaches the human central nervous system (24). In PD, a randomized, placebo-controlled trial of an oral resveratrol formulation, focused on safety and pharmacokinetics, has only recently been registered (NCT07592767). As an existing compound with a characterized safety profile, resveratrol is well suited to drug repurposing, which shortens the path from molecular finding to candidate therapy (25).

Our purpose in this study is to integrate transcriptomic datasets from postmortem brain tissue, iPSC– derived models, and established cell lines of PD. We propose an integrative framework designed to enable drug–gene interaction screening and thereby mitigate the heterogeneity inherent across PD models, ultimately facilitating the characterization of plausible therapeutic candidates.

## 2 Methods

### 2.1 Dataset selection and processing

Publicly available transcriptomic datasets were retrieved from the NCBI Gene Expression Omnibus (GEO, National Center for Biotechnology Information, Bethesda, MD, USA; www.ncbi.nlm.nih.gov/geo/). The database was screened between February 2025 and May 2026 using the search terms “Parkinson’s disease”, “postmortem”, “alpha-synuclein”, “dopaminergic neuron”, “LUHMES”, “SH-SY5Y”, and “iPSC”, restricted to *Homo sapiens*.

Datasets were included if they met all of the following criteria; (i) relevance to Parkinson’s disease, either as patient-derived material or as an established experimental PD model, (ii) clearly defined and separable disease and control groups, (iii) explicit description of the biological source of the samples (brain region, cell type or differentiation stage), (iv) human origin, (v) generation by RNA sequencing or microarray, with the platform and reference genome documented in the original submission, and (vi) availability of raw or processed expression data sufficient for independent reanalysis.

A total of 33 datasets meeting all inclusion criteria was identified and analyzed. Where a single GEO accession contained multiple independent comparisons (distinct brain regions, differentiation stages, culture conditions, doses or exposure durations), each comparison was processed as a separate dataset. These datasets were assigned to three model groups as postmortem human brain tissue (n = 5), iPSC-derived models (n = 16), and cell-based models (n = 12), with dataset characteristics including sample numbers per group, platform identifiers, and corresponding original publications (Table 1 and S1).

**Table 1.**
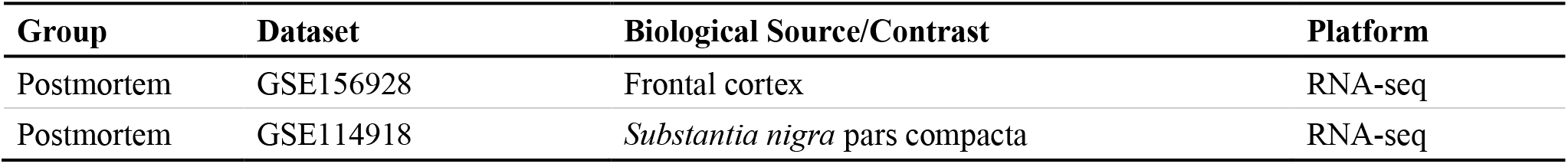
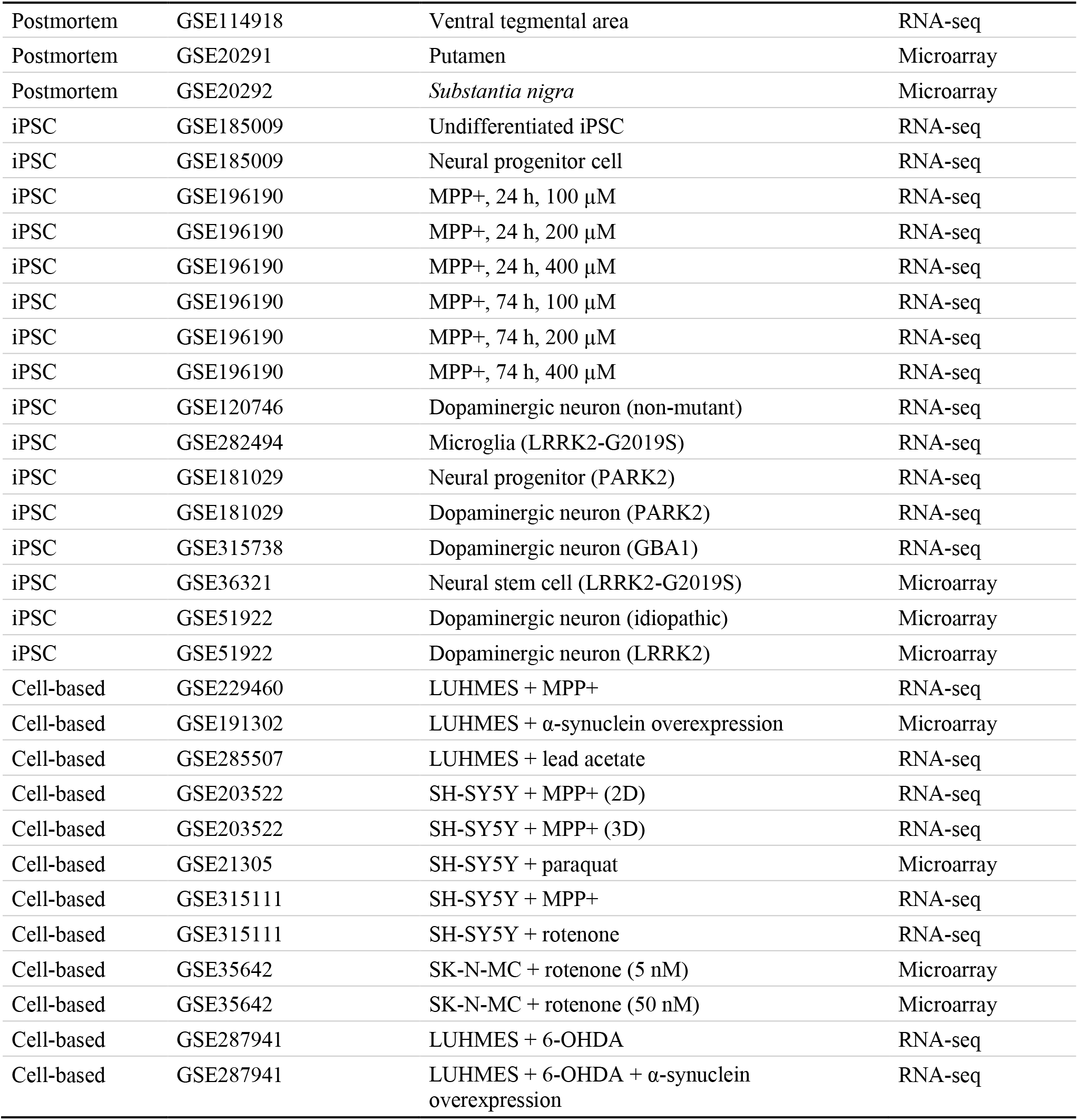
Transcriptomic datasets from different PD models for the analyses.

### 2.2 Transcriptomics for the PD models data

Raw RNA sequencing data in SRA format were retrieved and converted to FASTQ format using the Galaxy platform (Galaxy Project, Johns Hopkins University, Baltimore, MD, USA; usegalaxy.org). Adapter removal and quality trimming were performed with Trimmomatic (v0.39, Galaxy Version 0.39+galaxy2) using default adapter settings and a sliding-window approach across 4 bases with a minimum average Phred quality of 28 (SLIDINGWINDOW:4:28). Quality-filtered reads were aligned to the human reference genome GRCh38/hg38 using HISAT2 (v2.2.2, Galaxy Version 2.2.2+galaxy0) with default parameters. Gene-level counts were generated with featureCounts (v2.1.1, Galaxy Version 2.1.1+galaxy1) using the GRCh38/hg38 reference genome annotation, with feature type set to “exon” and strandedness set to unstranded (26, 27). Count matrices were imported into R (v4.3.3; R Foundation for Statistical Computing, Vienna, Austria). Genes with a total count below 10 across all samples were removed. Differential expression analysis was performed with DESeq2 (v1.42.1) on raw counts, comparing PD or treated samples against their matched controls. Variance-stabilizing transformation (VST) was applied for exploration, visualization, and co-expression analysis. Principal component analysis (prcomp) and sample-to-sample heatmaps (pheatmap v1.0.13) were used to assess sample clustering. Samples with visible separation from their group in both PCA and heatmap analyses were excluded before downstream analysis (26, 27).

Microarray datasets were retrieved with the GEOquery package (v2.70.0). Expression values were log2-transformed where required. Values were then quantile-normalized to correct for platform-specific variation using normalizeBetweenArrays from limma (v3.58.1). Where multiple probes mapped to a single gene symbol, expression values were averaged across probes. Probes without an annotated gene symbol were discarded, and, because differential expression was performed on this gene-level averaged matrix, each gene appears once by construction in the differential expression tables (28, 29). The sample clustering and outlier assessment were performed as described above. Differential expression analysis was carried out with limma (v3.58.1), fitting a linear model with disease status as the sole covariate and applying empirical Bayes moderation (28, 29).

### 2.3 Differential expression analysis for PD datasets

For both RNA-seq and microarray datasets, genes were considered differentially expressed when the Benjamini-Hochberg adjusted p-value was below 0.05 and |log2 fold change| exceeded 0.6. The same thresholds were applied across all 33 datasets to allow cross-dataset comparison. A threshold of 0.6 rather than 1.0 was chosen because effect sizes in postmortem brain tissue are typically modest, and a more stringent cut-off would have excluded biologically relevant changes in that model group (26–29).

### 2.4 Functional enrichment analysis

Gene Ontology (GO) and KEGG pathway enrichment analyses were performed with clusterProfiler (v4.10.1) on the combined list of significantly up- and downregulated genes from each dataset. GO annotations were obtained from org.Hs.eg.db (v3.18.0) and KEGG pathway data via KEGGREST (v1.42.0). The three GO ontologies (biological process, molecular function, and cellular component) were analyzed separately. The background gene universe was defined as all genes retained after filtering and tested for differential expression in the corresponding dataset. P-values were adjusted using the Benjamini-Hochberg method, and terms with both an adjusted p-value and a q-value below 0.05 were considered significantly enriched. The ten most significant terms from each ontology were visualized using enrichplot (v1.22.0) (26–29).

### 2.5 Protein-protein interaction network

Protein-protein interaction networks were constructed for each dataset from its significant DEG list using the STRING database (v12.5; STRING Consortium, University of Zurich, Switzerland, and EMBL, Heidelberg, Germany) via the REST API (string-db.org/api/json/network), accessed with httr (v1.4.7) and parsed with jsonlite (v2.0.0). Queries were restricted to Homo sapiens (NCBI taxonomy ID 9606) with a minimum required interaction score of 700. Gene symbols were submitted in batches of 50, and the interactions from all batches were merged into a single dataset-level network before topological analysis (27, 28). Network topology was analyzed with igraph (v2.1.4). Degree, normalized betweenness, normalized closeness, and eigenvector centrality were computed for every node, and genes were ranked by betweenness centrality. All genes present in the network, not only high-centrality nodes, were carried forward to drug-gene interaction analysis. Networks were visualized with ggraph (v2.2.1) (27, 28).

### 2.6 Drug-gene interaction analysis

All genes present in each dataset-level interaction network were queried against the Drug-Gene Interaction Database (DGIdb, Washington University in St. Louis, MO, USA; dgidb.org) between April and June 2026. Retrieved interactions were filtered to retain compounds annotated as “Approved” in the DGIdb regulatory approval field, and the reported interaction scores were recorded for each drug-gene pair (27, 28).

A panel of polyphenolic compounds was defined before the drug-gene interaction analysis. Compounds were selected based on documented antioxidant and anti-inflammatory activity, with priority given to those with published experimental evidence in PD models (30). The panel of polyphenols from PubChem comprised apigenin, baicalein, catechin, curcumin, epigallocatechin, epigallocatechin gallate, hesperidin, isoquercitrin, kaempferol, luteolin, myricetin, naringenin, quercetin, resveratrol, rutin, and troxerutin. Compound names in the DGIdb output were matched to the panel case-insensitively. For each recovered compound, the number of datasets in which it appeared was counted within each model group and across all 33 datasets, giving a frequency-based measure of cross-model consistency.

### 2.7 Correlation analysis through PD datasets

The relationship between expression profiles was examined in each dataset using Spearman’s rank correlation, computed with the cor.test() function in R on variance-stabilized (RNA-seq) or quantile-normalized log2 (microarray) expression values across all samples of a given dataset, including both disease and control groups. A correlation computed in this way reflects concordance in the direction of expression change between the corresponding genes across the disease-control contrast rather than co-regulation within a homogeneous population. The results were therefore interpreted as directional concordance. Correlation coefficients (ρ) and corresponding *p*-values were reported together with sample size for every dataset. Coefficients derived from fewer than six samples were considered exploratory and were reported without inferential interpretation.

### 2.8 Visualization and software

Drug-gene-dataset interaction networks were visualized as chord diagrams using Circlize (v0.4.16). Additional plots were generated with ggplot2 (v3.5.2) and EnhancedVolcano (v1.20.0). All analyses were conducted in R (v4.3.3, R Foundation for Statistical Computing) using RStudio (v2025.05.1 Build 513, Posit Software). Analysis scripts, including the marker gene panels, correlation workflow, and figure-generation code, were deposited on GitHub https://doi.org/10.5281/zenodo.22201827 and were cited in the Data availability section below.

## 3 Results and Discussion

### 3.1 The selection and processing of datasets

The study integrated 33 transcriptomic comparisons derived from 20 GEO datasets representing three complementary PD model systems: postmortem brain tissue (n = 5), iPSC-derived neuronal models (n = 16), and dopaminergic-like cell lines (n = 12) (Table 1). Twenty-four comparisons were generated using RNA sequencing and nine using microarray platforms, with cohort sizes ranging from 2 to 15 samples per condition (Supplementary Table S1). Collectively, the datasets captured four anatomically distinct brain regions, idiopathic and monogenic PD backgrounds (*LRRK2, PRKN,* and *GBA1*), and six neurotoxic or proteotoxic experimental paradigms, including a concentration- and time-dependent MPP⁺ treatment series. All datasets were processed using a standardized bioinformatics pipeline consisting of differential expression analysis, functional enrichment analysis, STRING network reconstruction, DGIdb-based drug-gene interaction screening, and *SIRT1-SNCA* directional concordance analysis.

### 3.2 Differential expression and functional enrichment

The number of differentially expressed genes (DEGs) varied substantially across comparisons, with a median of 2,121 DEGs and a range from 249 in GSE285507 to 10,978 in GSE229460 (Supplementary Table S1). *SIRT1* satisfied the DEG thresholds in 11 of 33 comparisons, showing upregulation in seven and downregulation in four. Similarly, *SNCA* was differentially expressed in 11 comparisons, including eight instances of upregulation and three of downregulation. Concurrent differential expression of both genes was observed in only four datasets (GSE156928, GSE315111 MPP⁺, GSE35642 5 nM, and GSE287941 6-OHDA+asyn).

Despite considerable transcriptomic heterogeneity across datasets, pathway enrichment analysis revealed a limited number of recurrent biological processes. MAPK signaling was the most consistently enriched KEGG pathway, reaching significance in 21 of 33 comparisons and representing the only pathway shared across all three experimental systems (postmortem: 4/5, iPSC: 9/16, cell-based: 8/12). p53 signaling (18/33) and cell-cycle pathways (14/33) were enriched primarily in iPSC-derived and cell-based models, whereas PI3K-Akt signaling (16/33) was particularly prominent in iPSC datasets (10/16). In contrast, PD-related pathways displayed a more restricted distribution. The KEGG terms ’Parkinson diseas’ and ’Pathways of neurodegeneration’ were each enriched in seven comparisons, comprising postmortem *substantia nigra* and ventral tegmental area samples together with five acute neurotoxin exposure models (MPP⁺, rotenone, and 6-OHDA). Oxidative phosphorylation was significantly enriched in five of these comparisons. Autophagy- and lysosome-related pathways were detected in nine datasets, most frequently in postmortem tissue (3/5) and toxin-exposed cell-based models (5/12) (Table S1).

### 3.3 Drug-gene screening: Resveratrol and SIRT1

Screening all network genes against DGIdb recovered approved interactions for eight of the panel polyphenols (Table 2 and S2, Figure S1). Resveratrol was one of the compounds recovered in all 33 comparisons and therefore in every model group, tissue, genetic background, and stressor, along with curcumin. Quercetin (28/33) and apigenin (22/33) followed, whereas the remaining four compounds (epigallocatechin g., rutin, catechin, hesperidin) were recovered in fewer than half of the comparisons. Resveratrol was linked to 30 distinct genes and was the only panel compound for which *SIRT1* was returned as a DGIdb target gene (Table 2 and S2, Figure S1). Resveratrol crosses the blood-brain barrier, a prerequisite for any CNS repurposing candidate, and its neuroprotective effect in PD models has been reported to depend on SIRT1 activity in SIRT1-inhibitor rescue experiments (23, 31).

**Table 2.** Polyphenolic compounds recovered by drug-gene interaction screening across the PD datasets.

| **Compound** | **Datasets (n/33)** | **Postmortem (n/5)** | **iPSC (n/16)** | **Cell-based (n/12)** |
|---|---|---|---|---|
| Curcumin | 33 | 5 | 16 | 12 |
| Resveratrol | 33 | 5 | 16 | 12 |
| Quercetin | 28 | 4 | 13 | 11 |
| Apigenin | 22 | 2 | 10 | 10 |
| Epigallocatechin g. | 13 | 2 | 7 | 4 |
| Rutin | 10 | 2 | 3 | 5 |
| Catechin | 9 | 1 | 2 | 6 |
| Hesperidin | 3 | 1 | 1 | 1 |

The most frequent genes that resveratrol was linked to were BRCA1 (13), BIRC5 (11), CCND1 (10), *SIRT1* (6), and ATAD5 (6) (Table 2 and S2, Figure S1). By function, the recurrent targets fell into three groups: cell cycle and DNA damage response (BRCA1, BIRC5, CCND1, CDK2, TP53, ATAD5), survival and stress signaling (PIK3CA, AKT1, PRKCA, FOXO3, HIF1A, SIRT1), and inflammatory signaling (NFKB1, NFKB2, RELA, IL1B, MAPK14). The cell-cycle genes were found almost exclusively in iPSC and cell-based comparisons (BRCA1 in 12 of 13, CCND1 in 10 of 10) and were predominantly downregulated (Figure 1). *SIRT1* showed the opposite distribution, with three of its six occurrences in postmortem brain (Figure 1A). The six comparisons in which *SIRT1* was linked to resveratrol spanned all three model groups: frontal cortex (GSE156928), SNc (GSE114918 SNc), putamen (GSE20291), *LRRK2*-G2019S neural stem cells (GSE36321), SK-N-MC cells exposed to 5 nM rotenone (GSE35642 5 nM), and LUHMES neurons exposed to 6-OHDA with α-synuclein overexpression (GSE287941 6-OHDA+asyn). *SIRT1* was upregulated in the frontal cortex, rotenone and 6-OHDA+α-synuclein comparisons, and downregulated in the SNc, putamen, and *LRRK2* comparisons (Figure 1B). *SIRT1* was also differentially expressed in five additional comparisons (GSE114918 VTA, GSE20292, GSE185009 iPSC, GSE315111 MPP⁺, GSE35642 50 nM) without linking to resveratrol.

**Figure 1.**
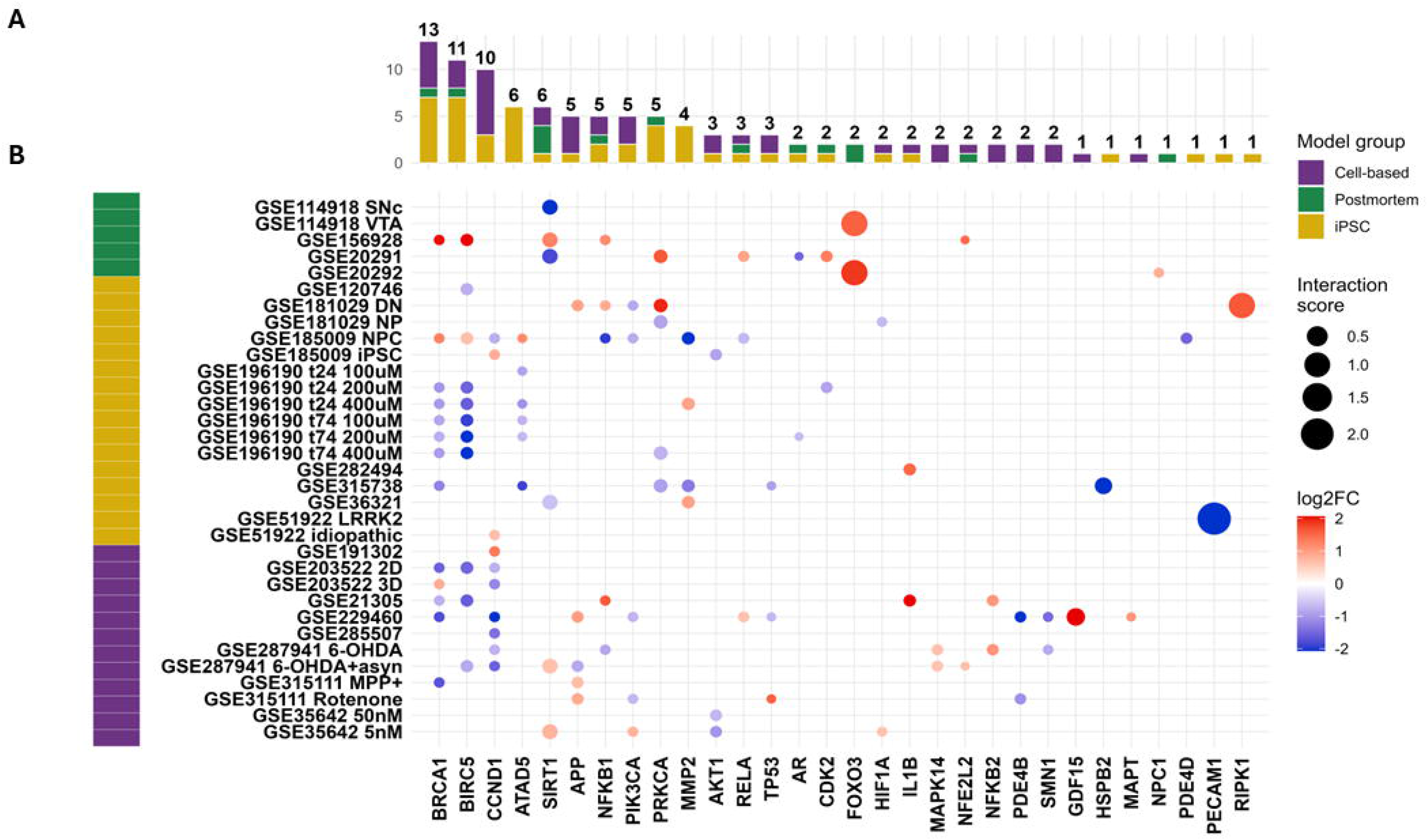
Resveratrol target genes across the 33 transcriptomic comparisons. **(A)** Number of comparisons in which each resveratrol target gene was recovered as a significantly differentially expressed gene, stacked by model group (postmortem, green; iPSC-derived, yellow; cell-based, purple); totals are printed above each bar. Genes are ordered by total frequency and share the x-axis with panel B. **(B)** Dot plot of the same gene set (x-axis) against all 33 comparisons (y-axis); the color strip on the left indicates the model group of each comparison. Only gene-comparison combinations in which the target met the differential expression criteria (adjusted p < 0.05 and |log2 fold change| > 0.6) are plotted; non-significant combinations are left blank. Dot color encodes log2 fold change on a diverging scale (blue, downregulated; white, unchanged; red, upregulated). Dot size encodes the DGIdb interaction score of the corresponding resveratrol-gene pair. A total of 30 distinct target genes was recovered overall.

### 3.4 SIRT1–SNCA directional concordance

To examine how *SIRT1* relates to α-synuclein, we compared the direction of change of *SIRT1* and *SNCA* in each comparison, using Spearman correlation across disease and control samples. Based on the sign of the correlation and which gene was differentially expressed, each comparison was assigned to one of four patterns (Figure 2). Assignments supported by a differentially expressed *SIRT1* were considered high confidence (10 comparisons), and those resting on *SNCA* alone were medium confidence (6) (Table S1). The two inverse patterns, marked by downward-pointing correlation arrows, contained all assigned postmortem comparisons (Figure 2). In the compensatory block, *SIRT1* bars were elevated, while *SNCA* values fell. This configuration was observed in the frontal cortex and *substantia nigra*, as well as in *PRKN* neural progenitors and 6-OHDA-treated LUHMES neurons. In the exhausted compensation block, the inverse relationship was retained, but the *SIRT1* bars pointed downward, most markedly in the SNc and VTA, and similarly in *LRRK2*-G2019S neural stem cells; in MPP⁺-treated LUHMES neurons, *SNCA* was upregulated instead.

**Figure 2.**
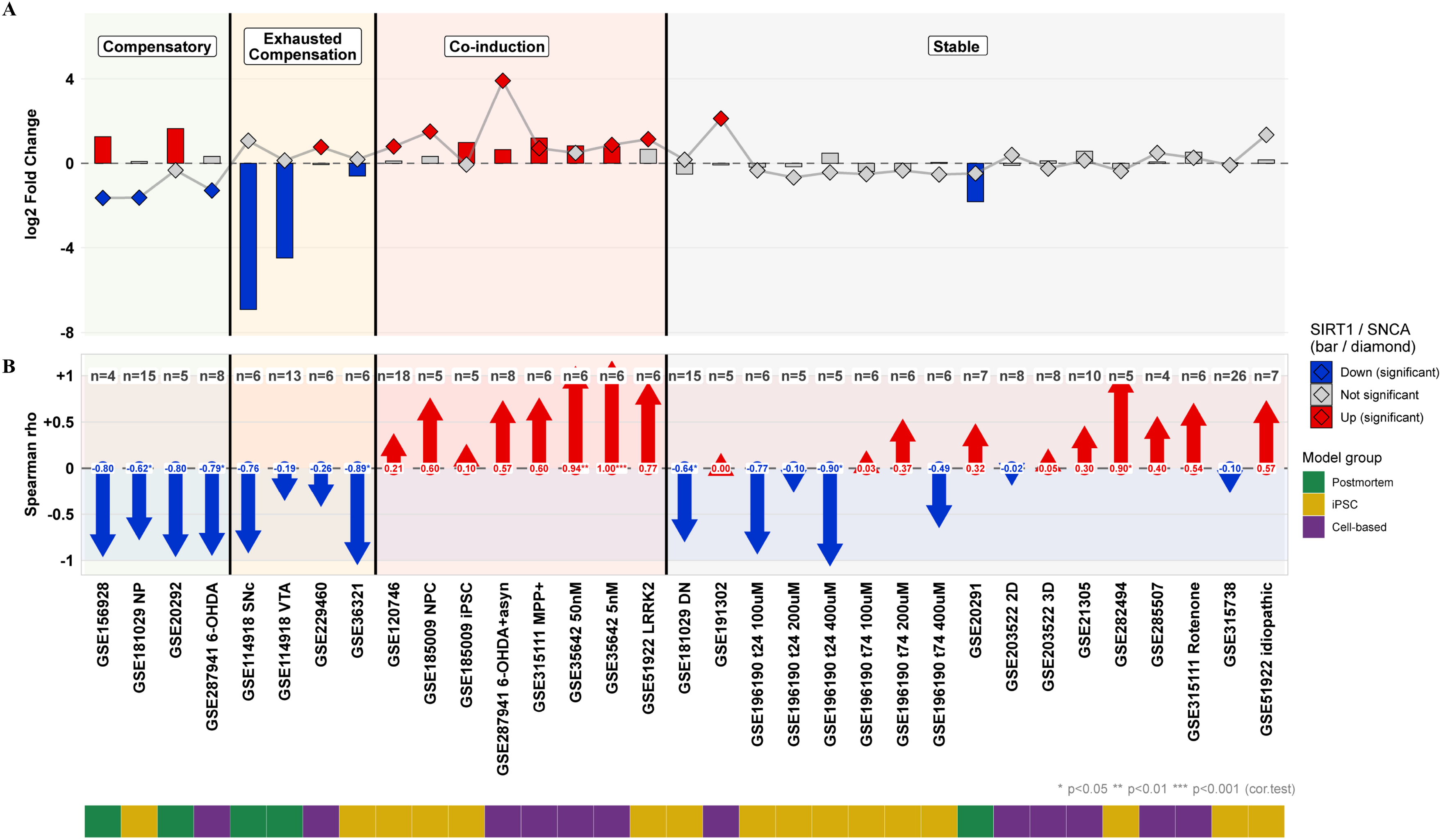
**SIRT1 and SNCA expression and directional concordance across the 33 comparisons, grouped by pattern**. Comparisons share a common x-axis across all three panels, and they are ordered into four pattern blocks (Compensatory, Exhausted Compensation, Acute Co-induction and Stable), separated by vertical lines and distinguished by background shading. **(A)** log2 fold change of SIRT1 (bars) and SNCA (diamonds joined by a grey line) was color-coded according to differential expression status: red, significantly upregulated; blue, significantly downregulated; grey, not significant. **(B)** Spearman rank correlation coefficient between SIRT1 and SNCA was computed with cor.test() and arrow direction and color-coded as the sign of the coefficient (red, positive; blue, negative); asterisks denote the coefficient’s own p-value ( p < 0.05, p < 0.01, * p < 0.001). Coefficients derived from fewer than six samples are displayed for completeness but are not interpreted individually.

The co-induction block, marked by upward correlation arrows, was dominated by toxin-exposed cell lines (Figure 2). Both genes were upregulated in SK-N-MC cells exposed to low-dose rotenone, SH-SY5Y cells exposed to MPP⁺, and LUHMES neurons exposed to 6-OHDA with α-synuclein overexpression, and *SIRT1* remained upregulated at the higher rotenone dose. The remaining members of this block were iPSC-derived comparisons without toxin exposure (undifferentiated iPSCs, neural progenitors, and dopaminergic neurons with or without the *LRRK2* mutation), in which the co-induction was driven mostly by *SNCA*.

### 3.5 Integration of resveratrol targets with SIRT1 and SNCA expression

To integrate the two parts of the analysis, the expression of *SIRT1* and *SNCA* was mapped onto the resveratrol target interactions for all 33 comparisons (Figure 3). The color of the *SIRT1* ribbons separated the model groups: significant decreases came mainly from postmortem brain, with the widest ribbons from the SNc and VTA, whereas significant increases came mainly from toxin-exposed cell lines. Ribbons from iPSC-derived models were largely non-significant for both genes. The widest *SNCA* ribbons originated from the two models engineered to overexpress α-synuclein (GSE287941 6-OHDA+asyn and GSE191302) and therefore reflect the experimental design rather than an endogenous response.

**Figure 3.**
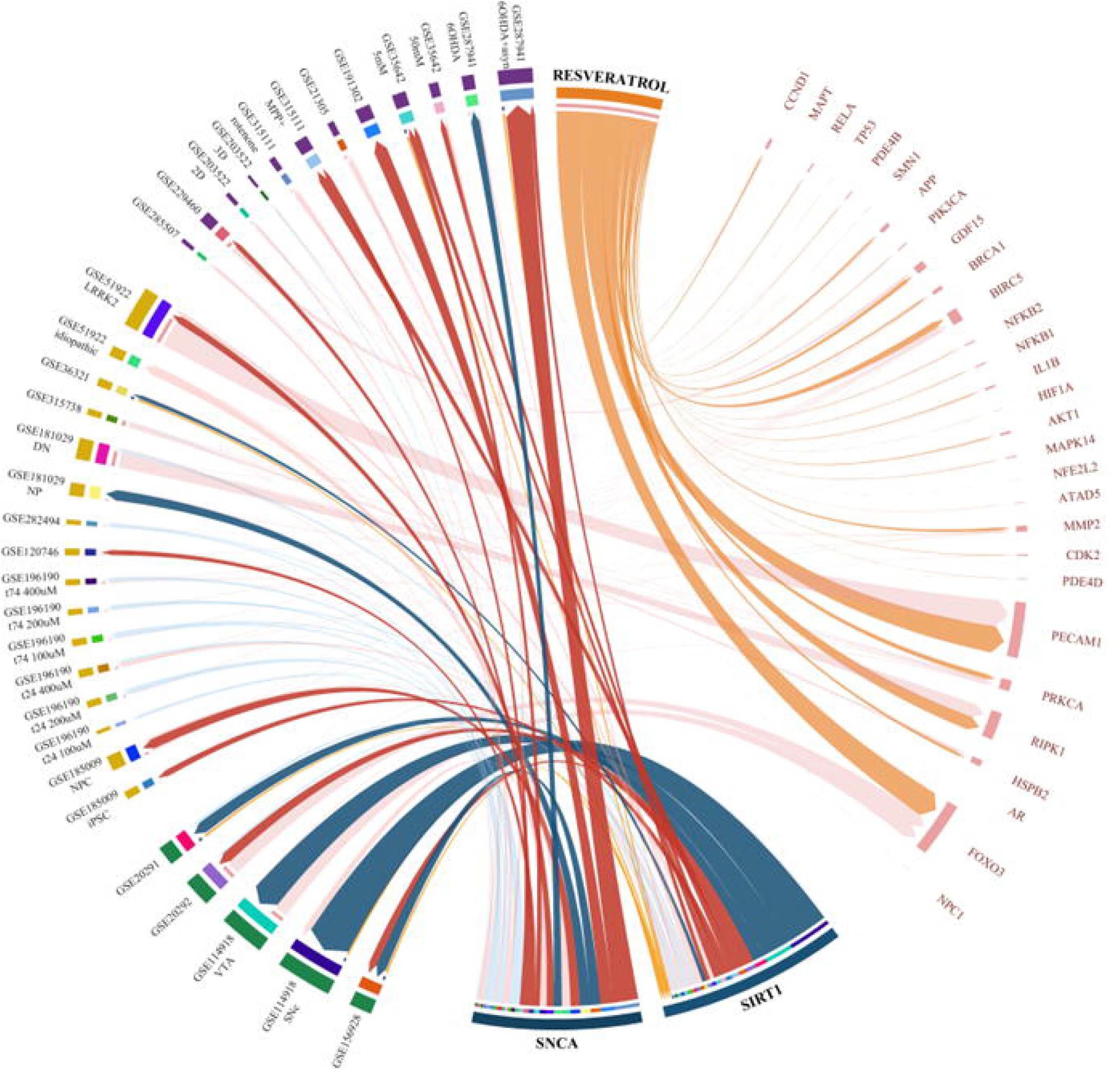
Chord diagram integrating resveratrol target interactions with SIRT1 and SNCA expression status across 33 Parkinson’s disease transcriptomic comparisons. Sectors on the left side of the circle represent the 33 transcriptomic comparisons, grouped and color-coded by experimental model system: postmortem brain tissue (green), iPSC-derived neurons (yellow), and cell-based models (purple). Sectors on the right represent resveratrol target genes retrieved from DGIdb, with SIRT1 and SNCA displayed as separate highlighted sectors. Orange ribbons connect Resveratrol to its target genes, weighted by interaction score; light pink ribbons connect each comparison to the target genes detected in that comparison, also weighted by interaction score. Ribbons connecting SIRT1 and SNCA to individual comparisons are weighted by mean |log2 fold change| and colored by expression direction and statistical significance: dark red indicates significant upregulation (padj < 0.05, log2FC ≥ 0.6), light red indicates non-significant upregulation, dark blue indicates significant downregulation (padj < 0.05, log2FC ≤ −0.6), and light blue indicates non-significant downregulation. Yellow-orange ribbons identify the six comparisons in which SIRT1 was itself returned as a direct resveratrol target by DGIdb, allowing these to be read against the SIRT1/SNCA expression pattern of the same comparison. SIRT1 was significantly upregulated in 7 and downregulated in 4 comparisons, while SNCA was significantly upregulated in 8 and downregulated in 3 comparisons.

The resveratrol side of the diagram revealed a trade-off between interaction strength and recurrence (Figure 3). The broadest target sectors, corresponding to the highest DGIdb interaction scores (PECAM1, FOXO3, RIPK1, GDF15 and HSPB2), each received ribbons from only one or two comparisons. Conversely, the cell-cycle genes BRCA1, BIRC5, CCND1 and ATAD5 received ribbons from many comparisons but formed narrow sectors. *SIRT1* was the only target that combined a relatively broad sector with ribbons from all three model groups. In postmortem comparisons, *SIRT1* co-occurred mainly with survival and antioxidant targets such as FOXO3, NFE2L2 and NPC1, whereas in cell lines it co-occurred with signaling targets such as PIK3CA, AKT1, HIF1A and MAPK14.

*SNCA* received no ribbon from resveratrol, so its link to resveratrol in this dataset runs entirely through *SIRT1* (Figure 3). The six highlighted comparisons in which *SIRT1* was a direct resveratrol target combined different *SIRT1*–*SNCA* configurations: *SIRT1* increased with *SNCA* decreased in the frontal cortex (Compensatory, Figure 2), *SIRT1* decreased in the SNc and *LRRK2*-G2019S neural stem cells (Exhausted Compensation, Figure 2), both genes increased in the rotenone and 6-OHDA+α-synuclein models (Co-induction, Figure 2), and *SIRT1* decreased without a matching *SNCA* change in the putamen (Stable, Figure 2). The resveratrol–*SIRT1* link was therefore present in both postmortem and experimental models and was not confined to a single state of the SIRT1–SNCA axis.

## 4 Conclusions

By integrating 33 transcriptomic comparisons from postmortem brain tissue, iPSC-derived models, and toxin-exposed cell lines, we demonstrated that a drug-gene screening approach converges on resveratrol, the only polyphenol identified across all comparisons and consistently linked to *SIRT1* (Table 1 and S1, Figure 1). *SIRT1* was among the few resveratrol targets shared between human brain tissue and experimental models, and its relationship with *SNCA* revealed distinct, context-dependent states such as an inverse association in postmortem tissue, a positive association under acute toxin exposure, and a stable relationship in iPSC-derived models (Figure 2).

The consistent identification of resveratrol across all 33 comparisons was not solely related to the number of its annotated targets, as quercetin possessed a larger target repertoire yet was recovered less frequently. Within the resveratrol interaction network, recurrent cell-cycle and DNA-repair genes, including *BRCA1*, *BIRC5*, and *ATAD5*, mainly reflected the proliferative biology of cultured cells (32, 33). In contrast, *SIRT1* combined a relatively high interaction score with consistent recurrence across all three model categories, with half of the observations originating from postmortem brain tissue (Figure 1). This finding underlines the importance of *SIRT1* at the intersection of experimental and human disease biology and identifies it as the resveratrol target with relatively strong translational relevance for PD.

The inverse *SIRT1-SNCA* relationship observed in postmortem tissue was consistent with the established role of *SIRT1* in promoting α-synuclein clearance through autophagy and FOXO-dependent signaling pathways (31, 34). In the compensatory state, increased *SIRT1* expression was accompanied by elevated *FOXO3* and *NFE2L2*, factors that cooperate with *SIRT1* to enhance antioxidant and autophagic defense mechanisms (Figure 2) (34, 35). This finding suggests the presence of an active neuroprotective response in selected brain regions, extending previous observations in PD and indicating potential variation across disease stages (14, 20, 21). By contrast, the exhausted compensation state, observed in the SNc and VTA where dopaminergic neurodegeneration is most pronounced (10, 13), may reflect a partial failure of this protective programme (Figure 2). Nevertheless, persistent *FOXO3* expression and ongoing autophagy-related signaling in the VTA suggest that downstream defense pathways remain at least partially functional (36). In *LRRK2*-G2019S neurons, where sirtuin activity is known to be impaired, *SIRT1* emerges as a particularly compelling therapeutic target (37). The coordinated induction of *SIRT1* and *SNCA* under acute toxin exposure appears to represent an early stress-responsive state, whereas the stable state observed in iPSC-derived models indicates that activation of the *SIRT1-SNCA* axis is selective rather than a universal consequence of PD-related perturbations (Figure 2 and 3) (38).

Collectively, these findings strengthen the role of resveratrol as a drug-repurposing candidate with an established safety profile, which penetrates the human central nervous system, and reduces α-synuclein accumulation in experimental PD models (23, 24). Moreover, highly improved bioavailability formulations have progressed to phase 2a clinical evaluation in PD (NCT07592767, 39, 40).

In conclusion, this cross-model analysis identifies *SIRT1* as the most consistent and translationally relevant resveratrol target in PD and demonstrates that its relationship with α-synuclein varies according to biological context (Figure 3). By linking a clinically advanced compound to a mechanistic axis conserved across both human brain tissue and experimental models, these findings provide a framework for identifying the disease stages and experimental settings in which resveratrol-mediated *SIRT1* activation is most likely to be beneficial, while supporting its continued development as a potential therapeutic strategy for PD.

## 6 Author contributions

F.H.: Conceptualization, Formal analysis, Investigation, Data curation, Visualization, Writing – original draft. E.S.: Conceptualization, Funding acquisition, Resources, Writing-original draft, review & editing. H.Ç.: Conceptualization, Supervision, Resources, Funding acquisition, Writing – original draft, review & editing. I.A.K.: Supervision, Resources, Funding acquisition, review & editing. All authors read and approved the final version.

## 8 Acknowledgements

The authors thank Simay Akpinar and Busra Bicak (CoRD-Omics Research Group at the Institute of Biotechnology, Gebze Technical University) for technical assistance in coding for the data analyses and drug screening, respectively.

## 9 Use of generative artificial intelligence

During the preparation of this study, the authors used generative artificial intelligence tools (ChatGPT, OpenAI; Claude, Anthropic) for two purposes: assistance in writing and debugging the R analysis scripts used for data processing and statistical analysis; and language editing, structural revision, and consistency checking of the manuscript text against the underlying analysis outputs. No generative AI tool was used to generate or interpret the primary transcriptomic data, to select datasets, or to formulate scientific conclusions. All AI-assisted code was reviewed, executed, and verified by the authors, and all reported values were checked against the raw analysis output. The authors take full responsibility for the content of this article.

## 10 Data availability

All datasets analyzed in this study are publicly available from the NCBI Gene Expression Omnibus (GEO) under the accession numbers listed in Table 1. Analysis scripts supporting the findings of this study are available at https://doi.org/10.5281/zenodo.22201827. The data that support the findings of this study, including Supplementary Tables S1 and S2, are available within the article and its Supplementary Information.

